# Neural Gene Network Constructor: A Neural Based Model for Reconstructing Gene Regulatory Network

**DOI:** 10.1101/842369

**Authors:** Zhang Zhang, Lifei Wang, Shuo Wang, Ruyi Tao, Jingshu Xiao, Muyun Mou, Jun Cai, Jiang Zhang

**Affiliations:** School of Systems Science, Beijing Normal University, Beijing, 100875, China; Key Laboratory of Genomic and Precision Medicine, Beijing Institute of Genomics, Chinese Academy of Sciences, Beijing 100101, China; University of Chinese Academy of Sciences, Beijing, 100049, China; Swarma Campus(Beijing)Technology Co., Ltd Beijing, 100035 China

## Abstract

Reconstructing gene regulatory networks (GRNs) and inferring the gene dynamics are important to understand the behavior and the fate of the normal and abnormal cells. Gene regulatory networks could be reconstructed by experimental methods or from gene expression data. Recent advances in Single Cell RNA sequencing technology and the computational method to reconstruct trajectory have generated huge scRNA-seq data tagged with additional time labels. Here, we present a deep learning model “Neural Gene Network Constructor” (NGNC), for inferring gene regulatory network and reconstructing the gene dynamics simultaneously from time series gene expression data. NGNC is a model-free heterogenous model, which can reconstruct any network structure and non-linear dynamics. It consists of two parts: a network generator which incorporating gumbel softmax technique to generate candidate network structure, and a dynamics learner which adopting multiple feedforward neural networks to predict the dynamics. We compare our model with other well-known frameworks on the data set generated by GeneNetWeaver, and achieve the state of the arts results both on network reconstruction and dynamics learning.

## 1 INTRODUCTION

Gene regulatory networks (GRNs) play a central role in the cell development and cellular identity. Transcription factors (TF) interact with each other and regulate hundreds to thousands downstream genes (JOHNSON et al., 2007), form the regulatory networks. Great efforts have been made in experimental method to wire this network together, which are crucial to decipher the basic mechanism of biology. By analyzing ENCODE data, a dense meta-network is constructed with multiple parts of the hierarchical networks exhibiting distinct properties (GERSTEIN et al., 2012). Meanwhile, the regulatory networks containing 762 human TFs interactions had been built. The authors use an integrative approach to systematically map combinatorial interactions among mammalian TFs (RAVASI et al., 2010).

Experimental method is not the only way to reconstruct regulatory networks. Alternatively, GRNs could be inferred from gene expression data. The conventional inference algorithms are based on mutual information (MARGOLIN et al., 2006), gene co-expression module (AIBAR et al., 2017) or Gaussian graphic model (TIAN; GU; MA, 2016). Most of the inference algorithms focus on the reconstruction of the regulatory networks from the un-ordered expression data but usually could not predict the dynamics of gene expression due to the lack of the time information of individual cell.

Single cell RNA sequencing (scRNA-seq) could measure gene expression levels in massive individual cells (ZHENG et al., 2017). Meantime, time label could be tagged on each cell computationally (QIU et al., 2017) or exprimentally (BRIGGS et al., 2018). Time-course scRNA-seq data has inspired new methods to infer both the network structure and the dynamics of gene expression (MATSUMOTO et al., 2017). However it over-simplifies the expression process because it is based on linear ordinary differential equations.

In this work, we present a purely data driven, model-free, heterogenous univesal deep learning model “Neural Gene Network Constructor” (NGNC) based on our previous Gumbel Graph Network (GGN) model (ZHANG et al., 2019). Here, heterogenous means each gene has a unique dynamic, and universal means any non-linear dynamic can be reconstructed by our model. The NGNC model can simultaneously infer the gene regulatory networks and reconstruct gene dynamics from time series gene expression data generated by GeneNetWeaver (GNW), which is referred as ‘golden standard’ for evaluation of network inference methods. Comparing to other network inference algorithms, our NGNC model could infer the structure of network with relative high accuracy. Meanwhile, the dynamic leaner in our model could accurately reproduce the dynamics of observed data with no restriction on the dynamic mechanism in advance.

## 2 METHODS

### 2.1 Model

The aim of the gene regulatory networks inference task is to reconstruct the regulatory network from time series gene expression data which could be measured as RNA-seq counts.

Following our previous work Gumbel Graph Network (GGN) (ZHANG et al., 2019), we designed NGNC with two modules: a network generator and a dynamic learner. The Network Generator module uses the Gumbel softmax trick (JANG; GU; POOLE, 2017) to generate the candidate adjacency matrix, which allows us to sample the adjacency matrix from the parameters in a differentiable way, and thereafter the stochastic gradient descent algorithm can be applied. Specifically, for an adjacency matrix *A* with *N* columns, we have a *N × N* parameterized matrix to optimize, with parameter *α*_*ij*_ denoting the probability that *A*_*ij*_ taking value 1. The candidate adjacency matrix can be sampled following the equation shown below:

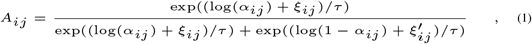

where *ξ*_*ij*_s and 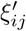s are random numbers following the gumbel distribution (NADARAJAH; KOTZ, 2004) and *τ* denotes the temperature. When *τ* is close to 0, the sampled *A*_*ij*_ will be close to 0 or 1.

The dynamic learner could predict the estimated values *X*_*t+1*_ for all variables(nodes) in the network at time *t* + 1 based on the candidate adjacency matrix generated by Network Generator and all observed real values *X*_*t*_ of all variables(nodes) measured at time *t*. In GGN model, we used a Graph Neural Network kernel to learn the homogeneous dynamics. but in gene regulatory networks, the regulation is heterogeneous for each gene since the TFs who regulate one single gene will combine together and interact in a non-linear way (RAVASI et al., 2010). So, we revise the dynamics learner to meet the need of heterogeneous regulation. In detail, the dynamics learner consists of several multi-layer perceptron (MLP), with each MLP corresponding to one single gene. The input of the MLP is a vector comes from the element-wised product of the gene expression vector *X*_*t*_ and the column vector *A*_*i*_ of adjacency matrix which representing the TF regulation acting on the corresponding gene *i*. The output of a MLP is 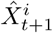, which is the estimated gene expression of gene *i* at time *t* + 1. The concatenation of all MLP outputs is the gene expression value 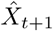 at time *t* + 1. We then can compare the output estimation 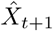 and the real expression value *X*_*t+1*_. Thus, the loss function of the whole system can be designed as:

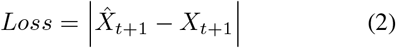

Then the stochastic gradient descent algorithm can be applied to minimize equation 2. The structure diagram of our model could be found in Fig 1.

**Figura 1:**
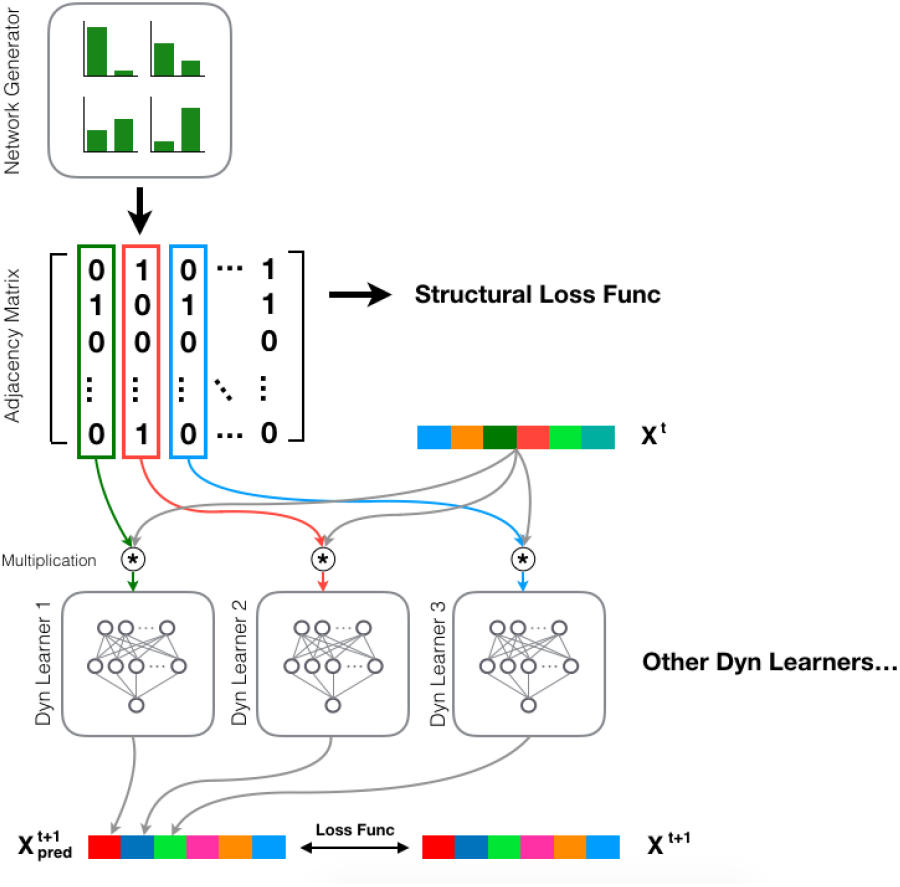
Architecture of NGNC: The NGNC model consists of two parts, a network generator and a group of dynamic learners. First, the adjacency matrix is generated by the network generator through Gumbel softmax sampling. Then, the element-wised products between gene expression vector *X*_*t*_ at time *t* and the column *i* of the adjacency matrix are calculated as the input for corresponding dynamic learner *i*. The dynamic learner *i*, which is a MLP, computes the output 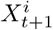, which is the estimation of the gene *i*’s expression at time *t* + 1. The concatenation of outputs of all dynamic learners is the estimation of all gene expression values *X*_*t+1*_ at time *t* + 1. The back-propagation process updates network generator and dynamic learners simultaneously.

On the testing process, we use ensemble learning technique to improve the prediction level. For each task, we choose three sets of randomly initialized parameters to train and let them vote for the reconstructed adjacency matrix. Other details about model and training process are described in SI.

## 3 RESULTS

### Time series data

The time series data is from the simulator GeneNetWeaver (GNW), which is used to evaluate different network inference methods systematically and comparatively (SCHAFFTER; MARBACH; FLOREANO, 2011). The time series are generated with default parameters in DREAM4_in-silico_Size-10 and DREAM4_in-silico_Size-100 configurations.

### Network inference

The Area Under Curve of ROC curve (AUROC) are used to evaluate the performance of the inference methods. We compare our method with other algorithms, such as Partial Correlation, Bayesian Network Inference and Mutual Information (see SI). As shown in Fig2, for simulated data with 10 TF and 100 TF, our model get higher AUC value than other algorithms. Especially for data with 100 TF, our model perform much better than other algorithms, indicating that our model could reconstruct the network with high accuracy even the network is large and complex.

**Figura 2:**
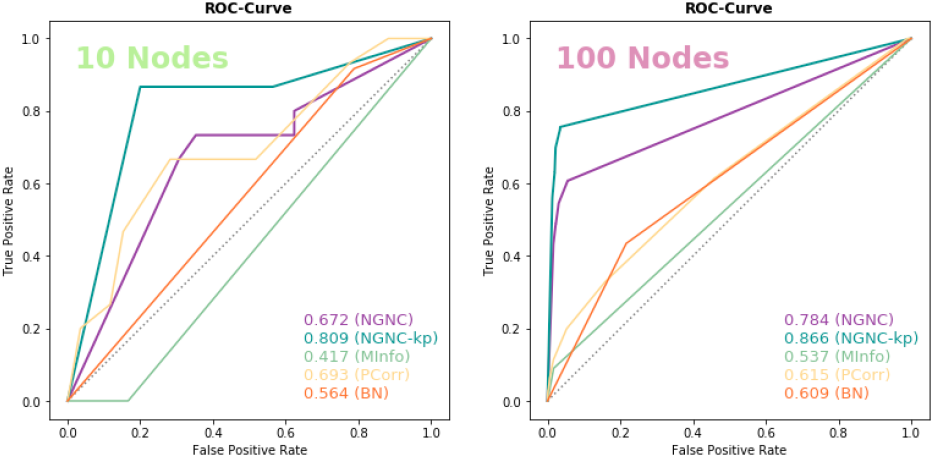
The performances (illustrated by ROC curves) of network inference methods: The ROC-Curve shows that the performance of NGNC (purple lines) is better than other network inference methods in the tasks with both 10 nodes and 100 nodes. Our NGNC could perform even better (Blue lines named NGNCkp) when some prior knowledge (see SI) is incorporated.

### Dynamic reconstruction

To test the performance of dynamic reconstruction, we fed nodes information at the first time step into the model and had it iterate 20 time steps. On 10 nodes and 100 nodes data, the mean absolute error (MAE) are 0.050 and 0.038. For further illustration, we draw both reconstructed dynamics and the observed time series on the same plot. As shown in Fig3, our reconstructed dynamics accurately reproduce the observed data. These results indicate that our method could learn the gene regulation dynamics correctly from the time series gene expression data. Meanwhile, the design of MLP dynamic learner in our model could approximate the underlying mechanism of gene regulation without any restriction on model of regulation such as linearality.

**Figura 3:**
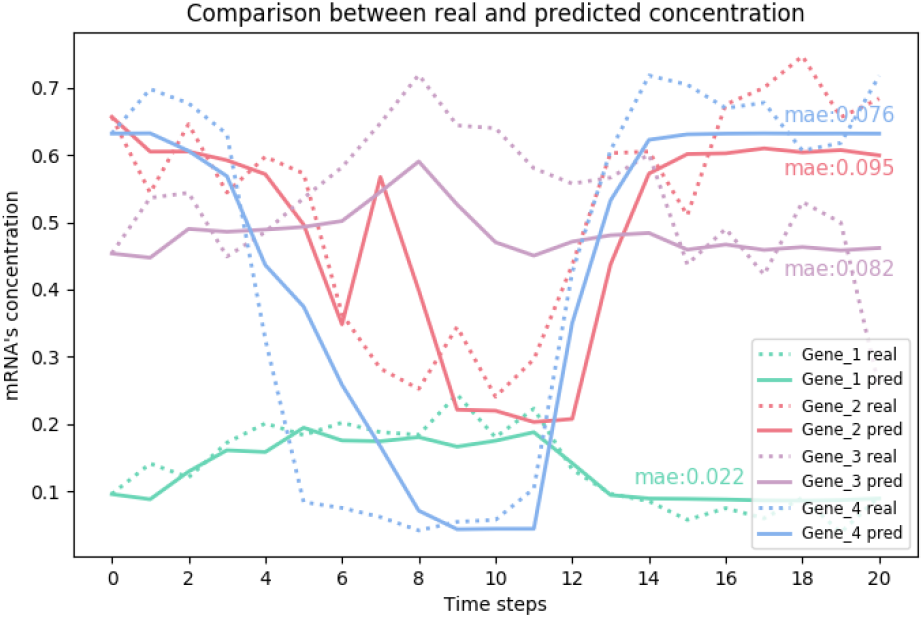
The comparison between the observed expression (real) and reconstructed dynamics (predicted) of several genes: In plots, solid lines represent predicted expression of genes and the dotted lines represent observed ones. One gene is depicted with one color. MAEs For each prediction curve are also plotted. As the plots show, the reconstructed dynamics faithfully reproduce the observed expression.

## 4 ACKNOWLEDGMENTS

This work was supported by grants from the National Natural Science Foundation of China [61673070 to J.Z. and 31571307 to C.J.]; the National Key RD Program of China [2018YFC0910402 to C.J.].

## REFERENCES

Aibar, S.; GonzÁLez-Blas, C. B.; Moerman, T.; Imrichova, H.; Hulselmans, G.; Rambow, F.; Marine, J.-C.; Geurts, P.; Aerts, J.; Oord, J. van den et al. Scenic: single-cell regulatory network inference and clustering. Nature methods, Nature Publishing Group, v. 14, n. 11, p. 1083, 2017.

Briggs, J. A.; Weinreb, C.; Wagner, D. E.; Megason, S.; Peshkin, L.; Kirschner, M. W.; Klein, A. M. The dynamics of gene expression in vertebrate embryogenesis at single-cell resolution. Science, American Association for the Advancement of Science, v. 360, n. 6392, p. eaar5780, 2018.

Gerstein, M. B.; Kundaje, A.; Hariharan, M.; Landt, S. G.; Yan, K.-K.; Cheng, C.; Mu, X. J.; Khurana, E.; Rozowsky, J.; Alexander, R. et al. Architecture of the human regulatory network derived from encode data. Nature, Nature Publishing Group, v. 489, n. 7414, p. 91, 2012.

Jang, E.; Gu, S.; Poole, B. Categorical reparameterization with gumbel-softmax. 2017.

Johnson, D. S.; Mortazavi, A.; Myers, R. M.; Wold, B. Genome-wide mapping of in vivo protein-dna interactions. Science, American Association for the Advancement of Science, v. 316, n. 5830, p. 1497–1502, 2007.

Margolin, A. A.; Nemenman, I.; Basso, K.; Wiggins, C.; Stolovitzky, G.; Favera, R. D.; Califano, A. Aracne: an algorithm for the reconstruction of gene regulatory networks in a mammalian cellular context. In: BIOMED CENTRAL. BMC bioinformatics. [S.l.], 2006. v. 7, n. 1, p. S7.

Matsumoto, H.; Kiryu, H.; Furusawa, C.; Ko, M. S.; Ko, S. B.; Gouda, N.; Hayashi, T.; Nikaido, I. Scode: an efficient regulatory network inference algorithm from single-cell rna-seq during differentiation. Bioinformatics, Oxford University Press, v. 33, n. 15, p. 2314–2321, 2017.

Nadarajah, S.; Kotz, S. The beta gumbel distribution. Mathematical Problems in engineering, Hindawi, v. 2004, n. 4, p. 323–332, 2004.

Qiu, X.; Mao, Q.; Tang, Y.; Wang, L.; Chawla, R.; Pliner, H. A.; Trapnell, C. Reversed graph embedding resolves complex single-cell trajectories. Nature methods, Nature Publishing Group, v. 14, n. 10, p. 979, 2017.

Ravasi, T.; Suzuki, H.; Cannistraci, C. V.; Katayama, S.; Bajic, V. B.; Tan, K.; Akalin, A.; Schmeier, S.; Kanamori-Katayama, M.; Bertin, N. et al. An atlas of combinatorial transcriptional regulation in mouse and man. Cell, Elsevier, v. 140, n. 5, p. 744–752, 2010.

Schaffter, T.; Marbach, D.; Floreano, D. Genenetweaver: in silico benchmark generation and performance profiling of network inference methods. Bioinformatics, Oxford University Press, v. 27, n. 16, p. 2263–2270, 2011.

Tian, D.; Gu, Q.; Ma, J. Identifying gene regulatory network rewiring using latent differential graphical models. Nucleic acids research, Oxford University Press, v. 44, n. 17, p. e140–e140, 2016.

Zhang, Z.; Zhao, Y.; Liu, j.; Wang Shuo, T. R.-Y. X. R.-Y.; Zhang, J. A general deep learning framework for network reconstruction and dynamics learning. arXiv preprint arXiv:1812.11482, 2019.

Zheng, G. X.; Terry, J. M.; Belgrader, P.; Ryvkin, P.; Bent, Z. W.; Wilson, R.; Ziraldo, S. B.; Wheeler, T. D.; Mcdermott, G. P.; Zhu, J. et al. Massively parallel digital transcriptional profiling of single cells. Nature communications, Nature Publishing Group, v. 8, p. 14049, 2017.

